# Enrichment of somatic mutations in schizophrenia brain targets prenatally active transcription factor bindings sites

**DOI:** 10.1101/2022.02.23.481681

**Authors:** Eduardo A. Maury, Attila Jones, Vladimir Seplyarskiy, Chaggai Rosenbluh, Taejong Bae, Yifan Wang, Alexej Abyzov, Sattar Khoshkoo, Yasmine Chahine, Brain Somatic Mosaicism Network, Peter J. Park, Schahram Akbarian, Eunjung Alice Lee, Shamil R. Sunyaev, Christopher A. Walsh, Andrew Chess

## Abstract

Schizophrenia (SCZ) is a complex neuropsychiatric disorder in which both germline genetic mutations and maternal factors, such as infection and immune activation, have been implicated, but how these two strikingly different mechanisms might converge on the same phenotype is unknown. During development, cells accumulate somatic, mosaic mutations in ways that can be shaped by the cellular environment or endogenous processes, but these early developmental mutational patterns have not been studied in SCZ. Here we analyzed deep (267x) whole-genome sequencing (WGS) of DNA from cerebral cortical neurons isolated from 61 SCZ and 25 control postmortem brains to capture mutations occurring before or during fetal neurogenesis. SCZ cases showed a >15% increase in genome-wide sSNV compared to controls (p < 2e-10). Remarkably, mosaic T>G mutations and CpG transversions (CpG>GpG or CpG>ApG) were 79- and 62-fold enriched, respectively, at transcription factor binding sites (TFBS) in SCZ, but not in controls. The pattern of T>G mutations resembles mutational processes in cancer attributed to oxidative damage that is sterically blocked from DNA repair by transcription factors (TFs) bound to damaged DNA. The CpG transversions similarly suggest unfinished DNA demethylation resulting in abasic sites that can also be blocked from repair by bound TFs. Allele frequency analysis suggests that both localized mutational spikes occur in the first trimester. We call this prenatal mutational process “*skiagenesis*” (from the Greek *skia*, meaning shadow), as these mutations occur in the shadow of bound TFs. Skiagenesis reflects as-yet unidentified prenatal factors and is associated with SCZ risk in a subset (∼13%) of cases. In turn, mutational disruption of key TFBS active in fetal brain is well positioned to create SCZ-specific gene dysregulation in concert with germline risk genes. *Skiagenesis* provides a fingerprint for exploring how epigenomic regulation and prenatal factors such as maternal infection or immune activation may shape the developmental mutational landscape of human brain.

## Introduction

While schizophrenia (SCZ) typically presents in adolescence or early adulthood, known causative factors appear to act during development. For example, germline mutations recurrently identified in SCZ, such as *NRXN1, TRIO* and others^1,2^, encode proteins implicated in synaptic development, while inherited non-coding variants conferring SCZ risk also tend to affect regulatory networks critical for brain development^3,4^. Furthermore, longstanding epidemiological evidence and animal model studies suggest that prenatal factors including maternal infection and immune activation in the first two trimesters^5–9^ can also increase SCZ risk, but how these nongenetic factors contribute to development of SCZ decades later remains unclear.

Somatic variants, present in only a fraction of cells in the body, occur throughout development^10–12^, and contribute to many neurodevelopmental conditions such as pediatric epilepsy^13^ and autism spectrum disorders (ASD)^14–17^. Analyzing the patterns and distribution of somatic variants has provided biological insight into cancer and neurodevelopmental disorders^10,13,18–25^, and revealed important molecular mechanisms that leave mutational footprints. We analyzed somatic mutations from deep WGS of DNA extracted from purified neurons from healthy individuals and individuals with SCZ to specifically examine mutations occurring during early prenatal development. Beyond an overall enrichment of somatic mutations in SCZ, we also observed two specific mutational footprints in SCZ cases that were absent in controls.

## Results

### Study design and variants discovery

We sequenced DNA from NeuN+ neurons of post-mortem dorsal lateral prefrontal cortex (DLPFC) samples from 61 SCZ individuals and 25 neurotypical controls (Figure 1A, Table S2, Methods) which identifies somatic mutations occurring prenatally, since post-mitotic neurons do not divide after development (Figure 1A). The brains were homogenized and nuclei stained for NeuN, and subjected to FACS using standard methods^26^. DNA was then extracted from 500,000-1,000,000 nuclei and sequenced without amplification (Methods). Median genome coverage was 267X, with no significant difference in coverage between cases and controls (Figure 1B). Mosaic somatic single nucleotide variants (sSNVs) were identified using best practices of the Brain Somatic Mosaicism Network (BSMN), which offers high sensitivity to identify sSNVs down to a variant allele fraction (VAF) of 0.02 in WGS^27^. The final call-set consisted of 3,284 sSNV calls (2,422 in SCZ and 862 in controls), with VAFs ranging from 0.02 to 0.38. Orthogonal validation with amplicon-based sequencing (Methods) confirmed 23/25 (92%), with VAFs highly correlated with WGS estimates (R-squared = 0.87, Figure 1C) suggesting a highly accurate call-set.

**Figure 1.**
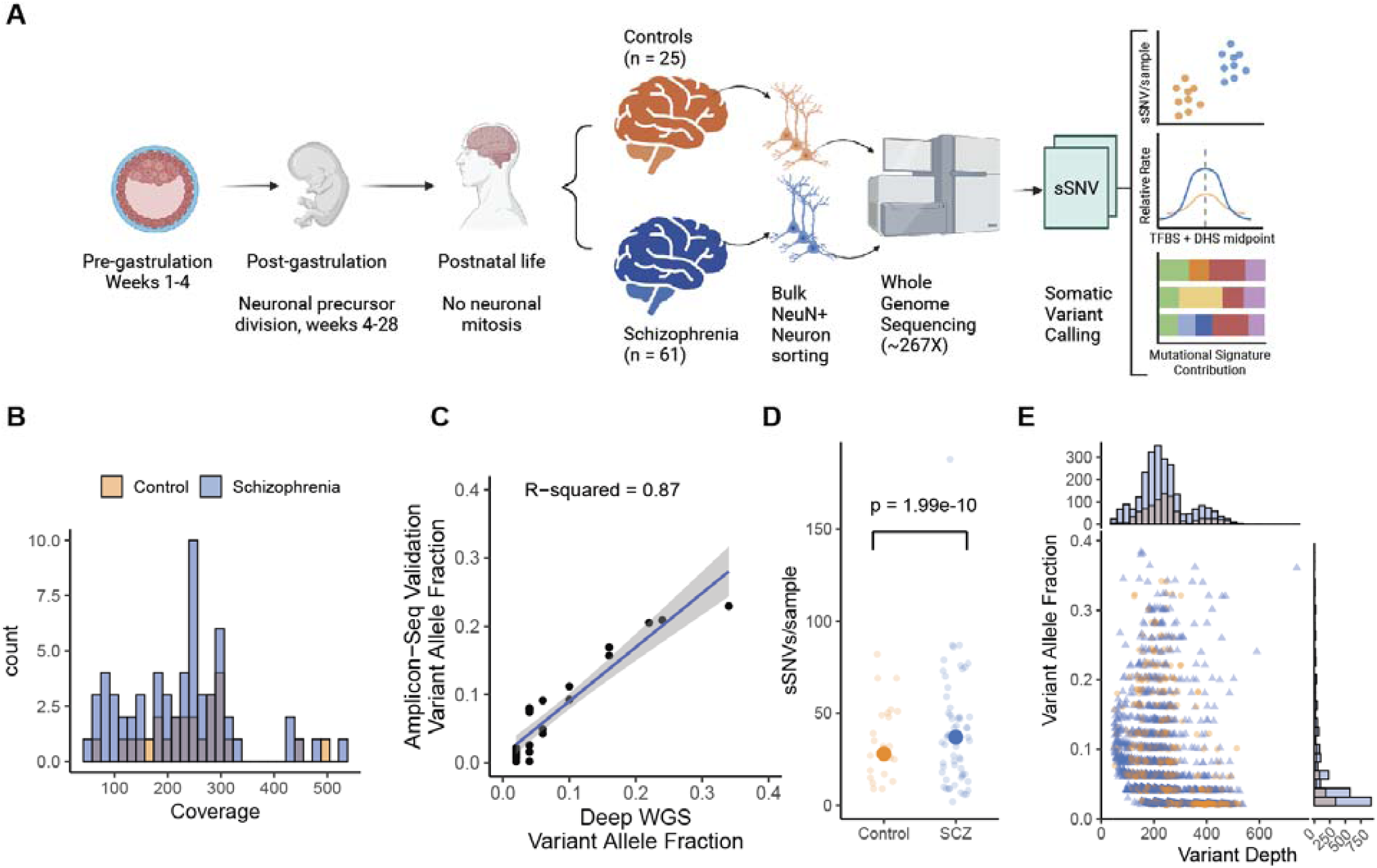
Increased genome-wide sSNV burden in schizophrenia cases compared to controls. A) Schematic of experimental and analysis design. Notably, neuronal clonal somatic mutations that are shared across neurons originate during prenatal brain development; occurring either before organogenesis (pre-gastrulation), resulting in somatic variants present in cells across multiple tissues, or during neuronal proliferation during neurogenesis. Mutations occurring postnatally in neurons are not clonal and hence undetectable with this method. B) Histogram of average sequencing coverage for schizophrenia cases and control samples. C) Scatter plot of Deep WGS variant allele fraction (VAF) for variant submitted to validation and the VAF from the validation amplicon sequencing. R-squared value best-fit line and 95% confidence band was computed from ordinary linear regression model. D) Scatter plot of number of sSNV per sample for schizophrenia cases and controls. Large points represent the sample means. The p-value was calculated using poisson regression. E) Scatter plot of VAF as a function of the number of reads covering the variant position. Margin plots show the corresponding marginal distributions for cases and controls for VAF and variant depth.

### Increased genome-wide sSNV burden in SCZ

Comparing the mean genome-wide sSNV counts revealed an unexpected increase in SCZ cases (39.7 per sample) compared to controls (34.5) (Figure 1D), a difference that was highly significant even after controlling for sample coverage, age of death, and standardized post-mortem interval (Poisson regression rate ratio (RR) = 1.30, 95% CI: [1.20-1.40], p=1.99e-10). One outlier SCZ sample with 188 mutations showed no technical or analytical anomalies that would justify its exclusion (see Supplemental Text), but the excess in genome-wide burden remained significant even after excluding this sample (p = 7.7e-5). VAF distributions were not significantly different between cases and controls (Kolmorogrov-Smirnov test, p=0.55, Figure 1E). Somatic copy number variant (sCNV) analysis revealed a somatic gain in one SCZ case, overlapping intron 1 and part of exon 2 of the *SORCS2* gene (Figure S1B, C), implicated in attention-deficit hyperactive disorder (ADHD), and bipolar disorder^28–31^, though a role of this gene in SCZ is not established (see Supplemental Text for more details). There was not a statistically significant enrichment of sSNV in GWAS loci previously associated with SCZ (Binomial regression, p = 0.936). Similarly, we did not detect somatic protein truncating variants (stop-gain, splice-site altering) or missense variants at genes implicated in SCZ in germline *de novo* or rare variant studies^1^ in our small sample size.

### Higher sSNV rate at active TFBS in SCZ

Analysis of the distribution of sSNV across the genome using fetal brain epigenomic tracks from Roadmap Epigenomes^32^, revealed an increased sSNV rate at open chromatin regions in SCZ neurons compared to controls. We observed higher sSNV rate in in SCZ compared to controls at DNase hypersensitivity sites (DHS) (binomial regression, p = 0.0008, Figure 2A), indicative of open chromatin. Conversely, we found a complementary lower sSNV rate in SCZ samples at H3K27me3 regions, associated with downregulation of genes and more closed chromatin^33^ (binomial regression, p = 0.0004, Figure 2A). On the other hand, we did not detect significant differences in sSNV rate at regions of increased fetal brain gene expression in cases compared to controls, nor a systemic transcriptional strand bias (Figure S2A), nor significant association between sSNV rate and replication time or replication fork direction (Figure S2B).

**Figure 2.**
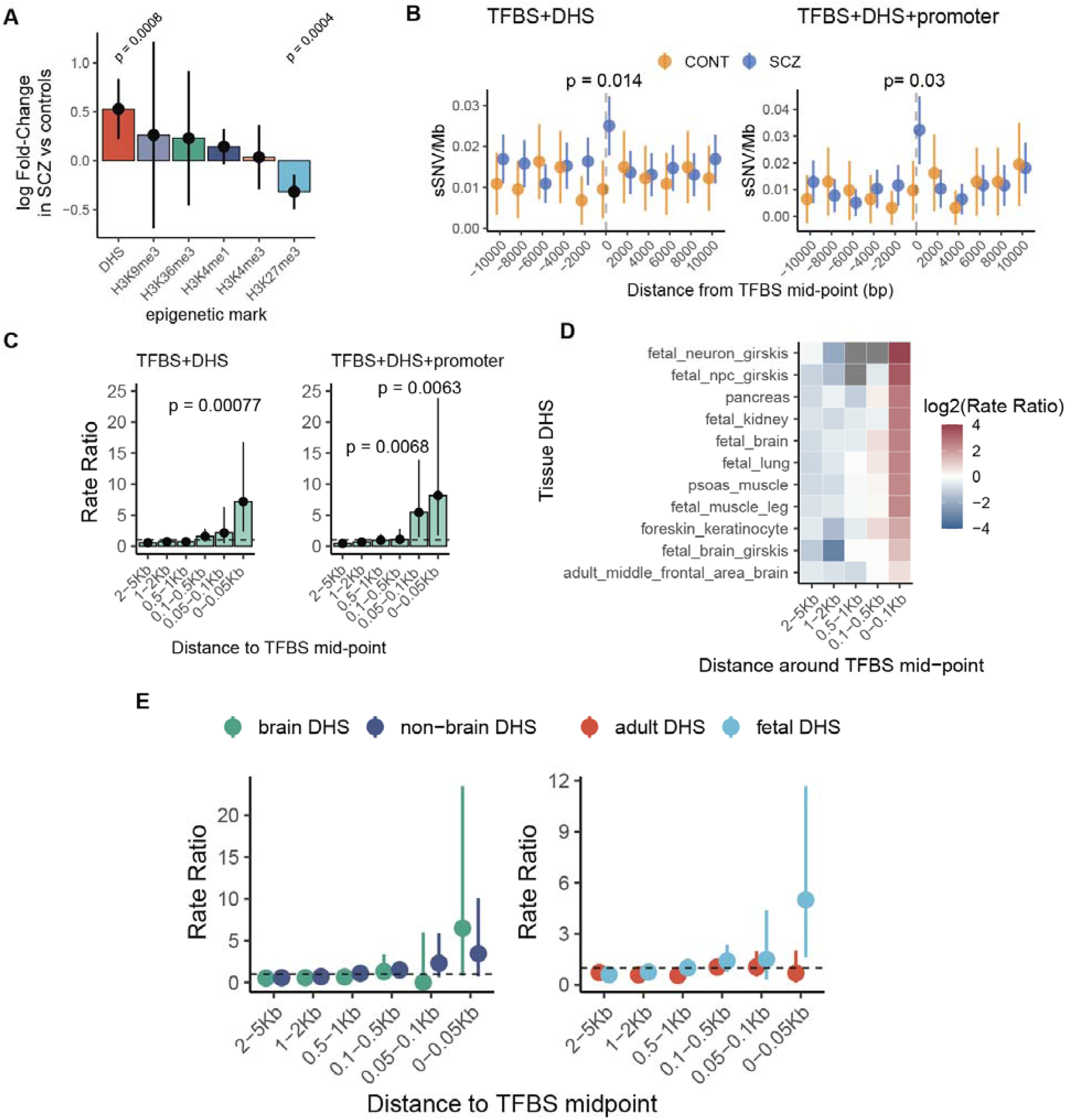
Increased sSNV rate at developmentally active transcription factor binding sites. A) Bar plot of binomial regression interaction term between epigenomic tracks and disease status. Positive values indicate enrichment in schizophrenia and negative values indicate depletion. Line ranges indicate 95% confidence intervals from binomial regression. B) Somatic SNV rate at +/-10Kb region from active transcription factor binding sites (TFBS) in fetal brain (TFBS+DHS), and active TFBS at promoter regions. C) Bar plot of rate ratios at binned regions around TFBS in schizophrenia. D) Heatmap of rate ratios in schizophrenia at TFBS using different DHS tracks. E) Rate ratio in schizophrenia around TFBS at DHS regions private to brain and non-brain regions (left), and private to adult or fetal tissues (right). For panels C, and D, p-values and confidence intervals were calculated using Poisson tests.

Previous studies in colorectal, skin, and other cancers observed an enrichment in point mutations at active TFBS, i.e. those overlapping DHS, due to hindrance of the DNA repair machinery by bound transcription factors (TFs)^34–37^. To test whether a similar phenomenon could explain the local increase in mutation rate at DHS regions in SCZ, we calculated the sSNV rates near the midpoint of active TFBS, accounting for the number of genomes sampled in each disease category (see Methods for details). We observed a robust increase in sSNV proximal (+/-1Kb) to the midpoint of the active TFBS in SCZ compared to controls (Poisson test, RR = 2.64 [1.18:6.93], p = 0.014, Figure 2B). This enrichment further increased at active TFBS overlapping promoter regions (Poisson test, RR = 3.32 [1.01:17.19], p = 0.03, Figure 2B). This enrichment over controls was observed across multiple TFs, thus the mutational process does not seem to be TF specific at this resolution (Table S3A, B).

Comparing the scale of the sSNV enrichment in SCZ near active TFBS to the expected genome-wide rate showed a 7.18-fold [2.33,16.8] enrichment at 50bp surrounding the TF mid-point (p = 7.69e-4, Figure 2C). This enrichment further increased to 8.48-fold [1.75:24.8] in promoter regions (p = 0.0057, Figure 2C). At distances exceeding 100bp from the TFBS midpoint, the mutational effect faded away completely; thus, this process is extremely localized. Significant mutational enrichment was seen at sites for many individual TFs essential for early embryonic, craniofacial, and/or nervous system development (Table S3C, D), with the highest nominal enrichment for Sp1^38^ (p = 5e-6), Rxra^39^ (p = 8e-5), Tfc12^40^ (p = 5e-4), and Fos/Jun^41^ (p = 1e-3).

### Prenatal TFBS are enriched for sSNV in SCZ

Comparison of sSNV rates at TFBS across DHS regions derived from different fetal and adult tissues suggests that the enrichment we observed in SCZ relates to fetal development. We observed significant enrichment of sSNV at active TFBS across all 11 tissues examined (including 7 fetal and 4 adult, Table S4, Figure 2D), suggesting that this mutational pattern is not obviously tissue specific (Figure 2D). and we observed similar sSNV increases at TFBS across DHS present in both brain-derived tissues and non-brain tissues (Figure 2E). On the other hand, sSNVs were enriched at TFBS overlapping private fetal DHS (RR=5.00, 95% CI [1.62-11.7], p = 3.67e-3], but not at TFBS overlapping private adult DHS (Figure 2E), consistent with the accumulation of these sSNV during prenatal development, yet potentially occurring in many fetal tissues rather than only in fetal brain.

### Specific sSNV patterns at TFBS

Analysis of specific base substitutions illuminated at least two potential sources of developmental mutagenesis. Measuring relative rates of sSNV across base changes at non-CpG sites, we observed a highly localized increase in sSNV rate of T>G substitutions (RR = 25.9, 95% CI [5.31:76.52], p = 2.4e-4) around 100 bp from the TFBS midpoint (Figure S3A), which was further enhanced at promoter regions (RR = 79.3, 95% CI[21.45:205.44], p = 2.69e-7, Figure 3A). T>G mutations made up a significantly higher proportion of sSNV in SCZ compared to control (Fisher Exact Test, OR= 2.19, 95% CI [1.58-3.09], p = 4.63e-7, Figure S4), while no other non-CpG base change was statistically significant (p > 0.05), suggesting a potentially disease-enriched mutational pattern.

**Figure 3.**
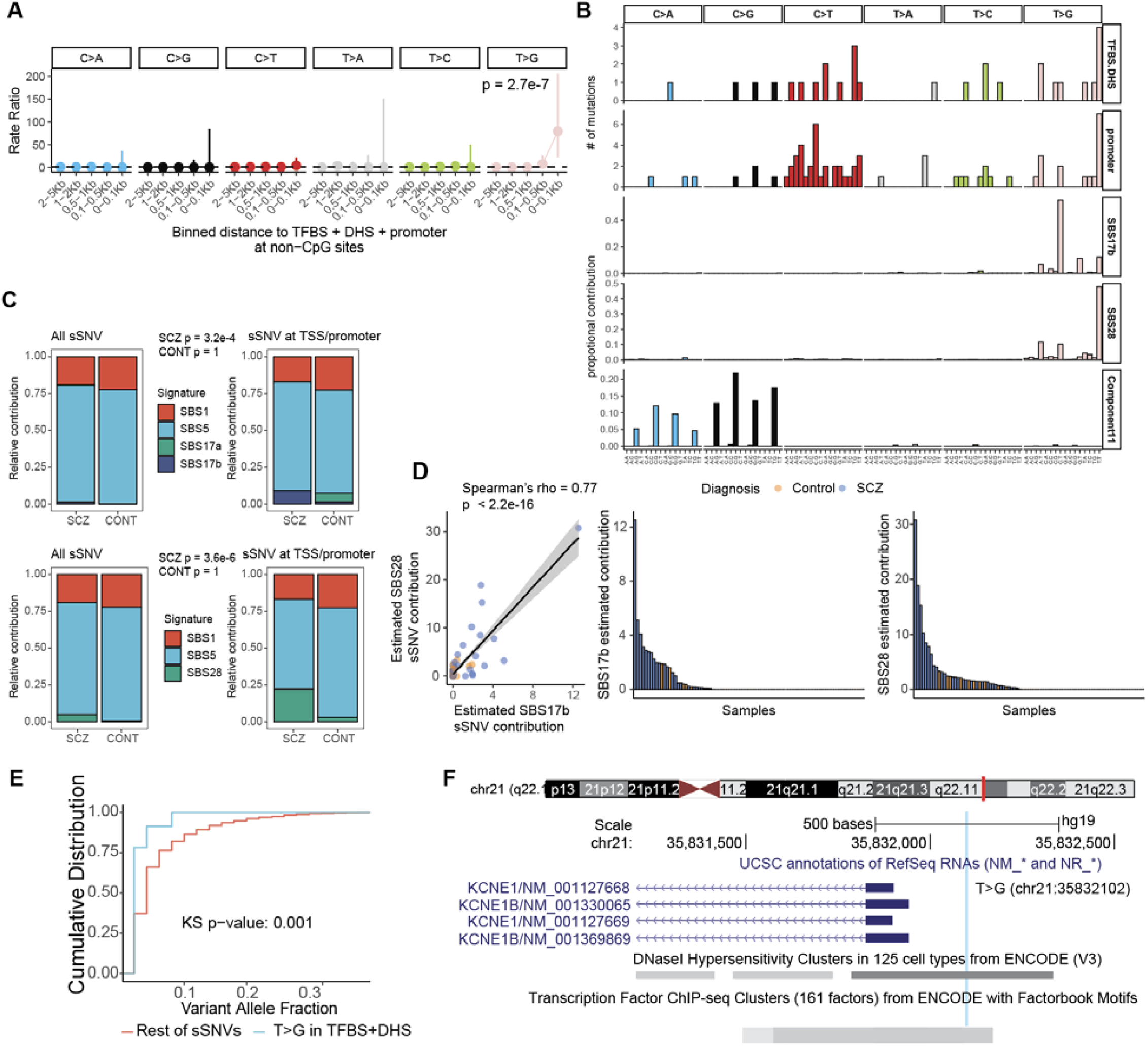
Increased somatic T>G base substitutions at active transcription factor binding sites in schizophrenia. A) Forest plots of rate ratios in schizophrenia of different base changes in active TFBS at promoter regions at non-CpG sites. P-values and 95% confidence intervals were computed using a Poisson test. B) Trinucleotide context plot of sSNV in schizophrenia at active TFBS and promoter sites, along with COSMIC signatures SBS17b, SBS18, and CpG transversion signature Component 11^44^. C) Proportion bar plots of the contribution of SBS17b, SBS28, along with SBS1 and SBS5, to the mutational spectrum in schizophrenia and control sSNV genome-wide and at promoter regions. P-values were calculated using a Fisher’s exact test. D) Contribution of SBS17b and SBS28 to the mutational spectrum in schizophrenia and controls samples as sSNV per sample. Left-plot: scatter plot of the contributions of SBS17b and SBS28. P-value was calculated using the spearman rank test. Middle and right plot show bar plots of the estimated sSNV contribution to each sample coming from SBS17b and SBS28. E) Empirical cumulative distribution plots of T>G mutations at active TFBS (blue) and the rest of sSNV in schizophrenia samples (red). P-value was calculated using Kolmorogrov-Smirnov test. F) Adapted Genome-Browser plot of a T>G mutations at chr21 in the promoter regions of the potassium channel gene *KCNE1*. Shades of gray on the DNAse Hypersensitivity tracks and TF Chip-seq annotations is proportional to the maximum signal strength observed in any cell line.

Previous studies in cancer showing enrichment of specific damage-induced sSNV at active TFBS have attributed them to steric interference of the TF with the DNA repair apparatus^34–37^, although other mechanisms have been proposed^35,42^. Somatic T>G mutations resembling COSMIC signature SBS17b^10^ at TFBS in gastrointestinal cancers have been attributed to oxidative damage^36^. Notably, the sSNV trinucleotide context at promoter sites and active TFBS in SCZ had a N[<u>T>G]</u>T mutational distribution resembling SBS17b (Figure 3B). We also observed a prominent peak at the T[<u>T>G]</u>T trinucleotide context, characteristic of SBS28^10^, which shares many features with SBS17b^10^. There was a significant increase in contributions of SBS17b and SBS 28 to sSNV mutations in SCZ (p=3.2e-4 and p = 3.6e-6, Figure 3C) at promoter sites compared to the expected genome rate, but not in controls (SBS17b and SBS28 p=1, Figure 3C). Brain samples from patients with ASD that were whole-genome sequenced at over 200X^17^ did not show this enrichment. These data suggest the intriguing possibility that prenatal oxidative damage, which does not get effectively repaired due to TF binding, might create mosaic sSNV at key TFBS, and in our data this phenomenon is relatively specific to SCZ.

Measuring the contribution of SBS17b and SBS28 across samples revealed that the T>G mutational processes was largely present in a subset of SCZ samples. As expected, the contribution of both signatures was highly correlated among samples (Spearman’s rho = 0.76, p < 2.2e-16, Figure 3D). While there was an overall significant enrichment of SBS17b (t-test, p=0.028) and SBS28 (t-test, p =0.034) in SCZ cases compared to controls, the enrichment in SCZ was mostly driven by a subset (∼8%) of SCZ cases with contributions of SBS17b or SBS28 >2 std above the mean (Figure 3D). Such a shared mutational process in a sizable group of individuals is most suggestive of a common maternal or fetal environmental process or exposure, though our data do not allow a definitive answer.

We observed that the VAF distribution of T>G mutations at active TFBS differed from that of non-T>G mutations outside of active TFBS (Kolmogorov-Smirnov test, p = 0.001, Figure 3E), allowing a few clues as to their timing. The VAF of T>G mutations at active TFBS was shifted toward lower values (0.02-0.08), suggesting that this process is not active during the first 1-3 cell divisions, but nonetheless begins during the first trimester^12^, since somatic mutations occurring after gastrulation are detected with very low sensitivity in 200X WGS^12,17^ (Figure 1A). However, given our low sensitivity for events with VAF lower than 0.02, it is possible that TFBS-associated accumulation of T>G mutations continues through later stages of fetal development as well.

Remarkably, we found 2 instances in which the exact same nucleotide substitution occurred at the exact same genomic position in two unrelated individuals with SCZ (Table S4), implying a remarkably narrow mutational target for this process. We did not observe any recurrent mutations in control samples or in review of other deep WGS samples^17^, and were able to validate these mutations by orthogonal methods (Methods, Table S4). Both recurrent sSNV were T>G/A>C mutations (Table S4). One occurred in the TFBS of ZNF343 within the promoter of the gene *KCNE1*, which encodes a potassium channel associated with congenital deafness and ventricular arrythmia^43^ (Figure 3F). The probability of observing 2 recurrent sSNVs by chance is exceedingly low (Poisson Test, p = 6.05e-6), suggesting that the mutational process driving these T>G mutations is highly localized in the genome.

A second potential mechanism was revealed by analysis of sSNVs at CpG sites, which showed up to 61.8-fold (95% CI [7.14:245.3], p=1.04e-4, Figure 4A, Figure S3B) enrichment of CpG>GpG in active TFBS at promoters compared to the expected C>G genome-wide rate. C>G and C>A transversions at CpG contexts are characteristic of a known mutational process (Component 11, Figure 3B)^44^ that reflects a footprint of enzymatic demethylation, which involves the resection of oxidated methyl-cytosine, creating an abasic site^45^ (Figure 4B). If the abasic site is not repaired before replication, CpG transversions occurs^46^. We speculate that this last step of demethylation could be similarly obstructed by TF binding, analogous to the interference between repair of lesions and TF binding^34,35,37^.

**Figure 4.**
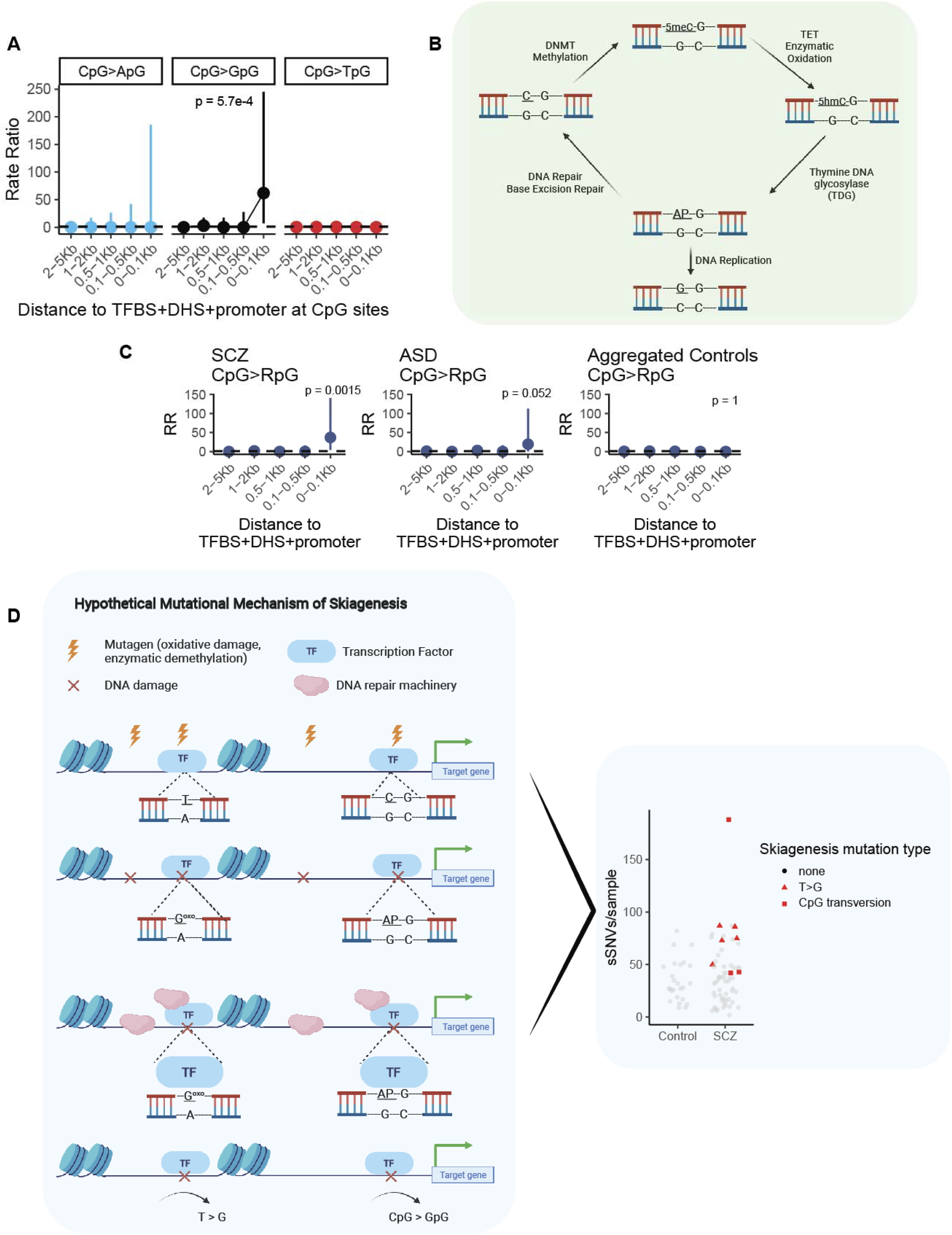
Increased somatic CpG transversions at active transcription factor binding sites in schizophrenia and hypothetical schematic of skiagenesis mutational model. A) Forest plots of rate ratios in schizophrenia of different base changes in active TFBS at CpG sites. B) Schematic of enzymatic demethylation mechanism resulting in CpG transversions. Abbreviations: 5meC, 5-methyl-cytosine; 5hmc, 5-hydroxymethyl-cytosine; AP abasic site. C) Forst plots of rate ratios of CpG transversions active TFBS in promoter regions from schizophrenia, Autism Spectrum Disorder, and aggregated control sSNV. For panels A and B p-values and 95% confidence intervals were computed using a Poisson test. D) Schematic of the model of skiagenesis. Abbreviations: TF, transcription factor; AP abasic site; G^oxo^, oxo-guanine. In the model, a mutational agent such as oxidative damage or enzymatic demethylation produces a lesion. The lesions can be repaired by DNA repair machinery, but these enzymes can be inefficient through steric inhibition by TF binding, resulting in mutation creation through replication. These mutations can then contribute to the higher sSNV burden observed in SCZ.

In addition to the enrichment of CpG transversions at active TFBS in SCZ (RR = 36.9, 95% CI [4.34:140.7], p=0.0015, Figure 4C), we observed a similar trend in bulk brain samples from individuals with ASD (RR=19.3, 95% CI [0.48:112.8], p=0.052, Figure 4C). This pattern was not observed in control neurons in the current dataset, nor in control bulk brain samples from the ASD study (Figure 4C). We similarly saw no enrichment of CpG transversions at close proximity to active TFBS in sSNVs obtained from a recent non-diseased twin study (Figure 4C)^47^. The enrichment of mosaic CpG transversions in disease cases (SCZ+ASD) compared to controls was statistically significant compared to aggregated sSNVs from non-diseased samples (Fisher’s Exact Test p = 0.004). The absence of acceleration of CpG transversion rate at active TFBS in controls again suggests that the interaction between TF binding and repair of abasic sites is specific to disease phenotype. By contrast we did not observe a significant difference in rate of somatic CpG transversions at CpG islands in disease compared to control (Fisher’s Exact Test p = 1, Figure S5), suggesting that CpG transversions in these latter regions might occur as part of normal development.

## Discussion

Our 267X WGS is sensitive to detect sSNVs shared by >2% of cells within a DNA sample^27^, indicating that the sSNVs detected in neuronal DNA from postmortem brain must have arisen in the dividing progenitor cells of those neurons, and hence be shared clonally among >2% of neurons (Figure 1A). Since neuronal progenitor proliferation ends by 24 weeks of gestation^48^, detected sSNVs must have occurred prior to that time (Figure 1A); the VAFs of specific variants, and their enrichment at chromatin regions open in many fetal organs, further implicate mutations arising in the first trimester, though potentially continuing later.

While our data is limited by the sample size and the number of mutations observed, they suggest mutational models that could explain the distinctive sSNV patterns observed in SCZ. One model consistent with the T>G process is that oxidative damage, as measured by the SBS17b/SBS28 mutational signatures, produce preferential sSNV accumulation at active TFBS due to hindrance of the DNA repair machinery by TFs bound to damaged DNA (Figure 4D). The genomic damage could reflect factors such as maternal infection or immune activation (MIA)^7^, which have been implicated in SCZ. Clinical epidemiology^5–7^ and animal model studies^7–9^ suggest that a portion of SCZ cases might be influenced by maternal infection, although the exact mutational footprint that this might create remains unknown; this suggests an area for further inquiry. Future studies with larger sample sizes might be needed to validate these findings using *de novo* signature extraction, and functional validation.

The mutational process involving CpG transversions at active TFBS could arise even earlier. We previously reported that CpG transversions make up ∼2.4% of all mosaic mutations from bulk brain tissue, potentially originating in the early zygote shortly after fertilization, when global DNA demethylation of the paternal and maternal genomes restores totipotency at the maternal-to-zygotic transition^44,45,49^. Alterations in this epigenetic process, either endogenous or exogenous, would predispose to somatic CpG transversions. The higher mean VAF of CpG transversions compared to T>G mutations at active TFBS (0.09 vs 0.02) is consistent with them occurring very early in development^12^. Comparison of mosaic mutation patterns between individuals with SCZ and controls is remarkable because the overall burden in CpG transversion is higher than seen in germline in both cases and controls^44^, but the effect of TF binding is unique to individuals with neuropsychiatric disease. While we favor a model in which these SCZ-specific processes may relate to known prenatally acting risk factors, other possibilities include SCZ-specific aspects of development, such as the division rate or TF binding time, that might also lead to TFBS being enriched with damage-induced mutations^50^. In any event, the T>G process and the CpG transversion process so far appear separate, observed in distinct individuals, and not so far shared in the same sample (Figure 4D).

We refer to these two mutational processes at TFBS as “skiagenesis”, from the Greek for “shadow,” because they occur in the shadow of TF binding. However, this term also captures the fact that despite the high risk they may create for SCZ, these sSNV are inaccessible to discovery by present-day clinical genetic testing. Their location in noncoding DNA and their exceedingly low VAF is only captured by deep WGS, which is expensive and informatically demanding.

Somatic SNV, enriched at essential TFBS active in development, are ideally suited to create risk for developmental brain dysfunction in SCZ. These mutations are predicted to be highly disruptive to the transcriptional regulation that underlies neuronal function, and may synergize with inherited and *de novo* germline SCZ risk genes that typically control gene dosage. Furthermore, the highly recurrent sites impacted by skiagenesis suggest how a nonspecific process can be channeled by specific TF binding to create recurrent patterns of mutation and recurrent risk for a behaviorally complex phenotype. Skiagenic mutations represent a fingerprint of prenatal factors that influence risk to SCZ and promise to help further dissect and classify these factors.

## Acknowledgements

E.A.M. is supported by the Harvard/MIT MD-PhD program (T32GM007753), the Biomedical Informatics and Data Science Training Program (T15LM007092), and the Ruth L. Kirschstein NRSA F31 Fellowship (F31MH124292). A.C. was supported by the NIMH grant (U01MH10681) and C.A.W. was supported by NIMH grant (U01MH106883, U01MH10681) through the BSMN. C.A.W is an Investigator of the Howard Hughes Medical Institute. C.A.W. and E.A.L. are supported by the Allen Discovery Center program, a Paul G. Allen Frontiers Group advised program of the Paul G. Allen Family Foundation. E.A.L. is supported by the NIH grants (K01 AG051791 and DP2 AG072437) and SUHF foundation. S.R.S. is supported by NIH grants R35GM127131, R01MH101244 and U01HG012009. Figures 1 and 4 were partly generated using Biorender.com. The authors thank Royce Park and Jennifer Wiseman and the flow cytometry core staff at the Icahn School of Medicine for technical support and the brain repositories associated with Common Mind Consortium for providing postmortem tissue.

## Contributions

E.A.M, A.J., V.S.B., A.C., S.A., C.A.W., S.R.S., C.R., and P.J.P. designed the study. E.A.M. and V.S.B designed and E.A.M. implemented the statistical methods and analysis with the help of V.B.S., and A.J.. E.A.M. performed burden analyses and mutational signature analyses. A.J. performed mutation calling with assistance of T.B., A.A., and P.J.P.. A.J. annotated the calls. C.R. and S.A. performed cell sorting and sequencing experiments. Y.W., T.B., and A.A. performed the sCNV calling. S.K. and Y.C. performed validation experiments. E.A.M., V.B.S., A.J., E.A.L, S.R.S., A.C. and C.A.W. wrote the manuscript.

## Methods

### Sample preparation and sequencing

Frozen post-mortem DLPFC (dorsolateral pre-frontal cortex) pulverized samples of subjects (61 schizophrenic and 25 control) were obtained from the Mount Sinai Brain Bank, part of the NIH NeuroBioBank. All specimens were deidentified, and all research was approved by the CommonMind Consortium. No statistical methods were applied to predetermine sample sizes, and rather we attempted to obtain data from all the affected and control frozen brains available to us at the time of the study and within the budget constraints of the project. Data collection and batching of samples were not randomized. We isolated NeuN+ (Anti-NeuN-Alexa488 (Cat# MAB377X, EMD Millipore) antibody) nuclei from DLPFC tissue samples using fluorescence-activated nuclei sorting^26^, followed by standard proteinase-K based DNA isolation with phenol-chloroform cleanup and ethanol precipitation. Sequencing libraries were then prepared with the Illumina TruSeq DNA PCR-free kit, according to the manufacturer’s standard protocol (350bp fragment design). We quantified sequencing libraries using the KAPA Library Quantification Kit (a real-time PCR methodology), and libraries were sequenced at the GeneWiz sequencing facility (NJ, USA) on an Illumina HiSeq X Ten platform, to yield 150bp paired-end reads. Sequencing experiments aimed for a minimum yield of 200x coverage per sample, and the average coverage obtained across all samples was 267x.

### Somatic SNV calling and filtering

Somatic SNVs were identified from WGS sequencing data using the best practices workflow from the Brain Somatic Mosaicism Network ^27^. Briefly, fastq files were aligned to the GRCh37 reference genome using bwa v0.7.17 ^51^, and preprocessed using the GATK best practices. Raw variants were then called using GATK Haplotypecaller ^52^ using a ploidy that corresponds to 20% of the overall sequencing coverage (i.e. ploidy of 50). Variants were then filtered if they fell on genomic regions labeled by 1000 Genomes Strict Mask ^27^. Variants with a GnomAD ^53^ population allele frequency >0.001 were filtered as well as variants with variant allele frequencies close to 0.5 (binomial test p < 1e-6) to remove potential germline variants. Candidate sSNVs were required to have >4 independent non-duplicated supporting reads with mapping quality of 20. A panel of normals filter from the 1000 Genomes Project was also used to remove variants that might occur from technical artifacts. The pipeline is readily accessible along with instructions at https://github.com/bsmn/bsmn-pipeline, which was run using the AWS ParallelCluster (https://github.com/aws/aws-parallelcluster) with the following configuration settings (https://github.com/bintriz/bsmn-aws-setup).

### Somatic copy number variant calling

We performed somatic CNV analysis on samples with 75 samples with mean coverage larger than 100x. We excluded 19 samples (MSSM_033, MSSM_063, MSSM_065, MSSM_069, MSSM_116, MSSM_118, MSSM_158, MSSM_192, MSSM_201, MSSM_266, MSSM_287, MSSM_291, MSSM_293, MSSM_299, MSSM_308, MSSM_309, MSSM_310, MSSM_331, MSSM_338) with coverage less than 100x. In addition, we excluded MSSM_164 with noisy signals in both read depth and allele frequency. Candidates for somatic CNVs were generated by CNVpytor ^54^ with the caller gathering information from both read depth and split in B-allele frequency of germline SNPs called using GATK haplotype caller run with ploidy=2. Analysis was conducted with two bin sizes: 100 kbps and 10 kbps.

We then applied filters to exclude false positive candidates and germline CNVs. We considered as false positives the following: 1) calls with adjusted p-value from CNVpytor larger than 0.05/(number of samples*3*109/bin size); 2) calls with <50% of well mapped bases (P-bases) as defined by the 1000 Genomes Project; 3) calls with >5% of non-sequenced reference (N-bases); 4) calls only supported by read depth (p-value from BAF signal > 0.01) and with predicted cell frequency <5%; 5) calls with predicted cell frequency <10%; 6) calls found in multiple samples (two calls are considered the same if overlap by 50% reciprocally). We additionally filtered out calls with length <= 3 of bins due to the boundary effects, which may lead to underestimation of cell frequency for germline CNVs.

We were not able to resolve breakpoints for the somatic duplication. Thus, we imputed two haplotypes by phasing germlines SNPs using population haplotypes and then confirmed that the frequencies of the two haplotypes were different.

### Amplicon Validation

Custom primers were designed for each candidate variant using the default settings in Primer3^55,56^ to generate 150-300bp amplicons. The primers were commercially synthesized (IDT) and tested on human genomic DNA (Promega) to confirm generation of only one amplicon product at the expected size. Then 10-50ng of genomic DNA from patients (based on sample availability) were used to create amplicons for sequencing, purified using 2X AMPure XP, and run on a gel for quality control. Amplicons from different samples were pooled together and illumina sequenced to achieve at least 10,000 reads per each unique amplicon. The raw reads were aligned to the reference genome (hg19) and the visualized on Integrative Genomics Viewer (IGV) to confirm the presence of each candidate variant. The variant allele frequencies were calculated based on the total number of REF and ALT alleles.

### Variant Annotation

For schizophrenia GWAS loci we used Table S4 of Pardinas2018 (DOI:10.1038/s41588-018-0059-2).

### Epigenomic mark enrichment

To test the for enrichment of epigenomic tracks (H3K27me3, DHS, H3K4me1, H3L36me3, H3K9me3, H3K4me3) in SCZ cases compared to controls we modeled the number of mutations *Y* at each track region *i* as a binomial outcome, such that:

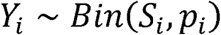

where *S* is the number of sites available to be mutated, and *p* is the probability of a site being mutated. For each track we constructed a matrix with N, the number of regions, times 2 rows (one for each disease category) and 3 columns (for the intercept, track signal, and diagnosis). So that we can estimate the relationship between each track’s signal and diagnosis status as a log binomial regression:

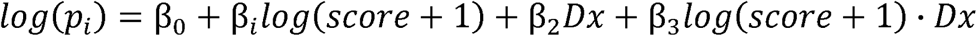

where score is the signal of each track respectively, and *Dx* is the diagnosis status. We considered a result significant if β_3_ ≠ 0, which we interpret as the excess effect of the epigenetic mark on somatic mutation rate in SCZ cases compared to controls. We used the *glm* R package to estimate these parameters. The broadpeak tracks were obtained from Roadmap Epigenomics from sample E081^32^.

### Transcription factor binding site track

We aggregated the hg19 TFBS bed files from Vorontosov *et al*^57^ using transcription factor tracks with highest reliability and experimental and technical reproducibility (A tracks). Since these tracks are an aggregation across experimental designs, they represent TFBS that are not necessarily tissue specific.

### DNAse hypersensitivity tracks

We obtained the DHS tracks from ENCODE^58^ and Roadmap Epigenome^32^. We also obtained tracks from fetal neuron, neuro-progenitor cells, and fetal brain from Girskis *et al*.^59^ For a complete list of the tracks and how to access them see Table S5. For most of the analyses involving DHS we used the broad peak calls with an FDR of 0.01 of fetal brain from sample E081 from Roadmap Epigenome^32^, unless otherwise stated.

### Comparison of sSNV rates between cases and controls at active TFBS

We compared the sSNV rates per Mb in a range of +/-10Kb from the TFBS mid-point. For this analysis the TFBS bed file was filtered by overlaps with the top 10% DHS regions from fetal brain (Table S5) and promoter regions. The promoter regions were defined as 2.5Kb upstream from transcription start sites as defined by *Ensembl* transcripts. The 20Kb range was binned into ∼2Kb windows and a Poisson test was used to compare the rates in SCZ and control mutations, using the *genomation* R package^60^. We adjusted by the number of samples in each disease category by multiplying the number of sites covered on each bean by the number of cases and controls respectively.

### Comparison of sSNV rates with genome-wide rates

We compared the sSNV rates per base pair at different distance intervals from the TFBS mid-point. For this analysis the TFBS bed file was filtered by overlaps with the top 5% DHS regions from fetal brain (Table S5) and promoter regions. The promoter regions were defined as 2.5Kb upstream from transcription start sites as defined by *Ensembl* transcripts. The number of mutations from the next interval closest to the TFBS midpoint was subtracted from the subsequent interval to make each interval independent. We used a Poisson test to compare the sSNV rate at each interval, using the genome-wide rate as the expected rate.

### Signature contribution estimation

We estimated the fraction mutational signatures SBS17b, SBS28, SBS1, and SBS5 from sSNV in cases and controls with the *MutationalPatterns* R package^61^. We estimated these fractions across all sSNV and at promoter regions defined as 2.5Kb upstream from transcription start sites as defined by *Ensembl* transcripts. The latter was used since it provided sufficient number of mutations for reliable signature estimation compared to active TFBS.

## Supplementary Figures

**Figure S1.**
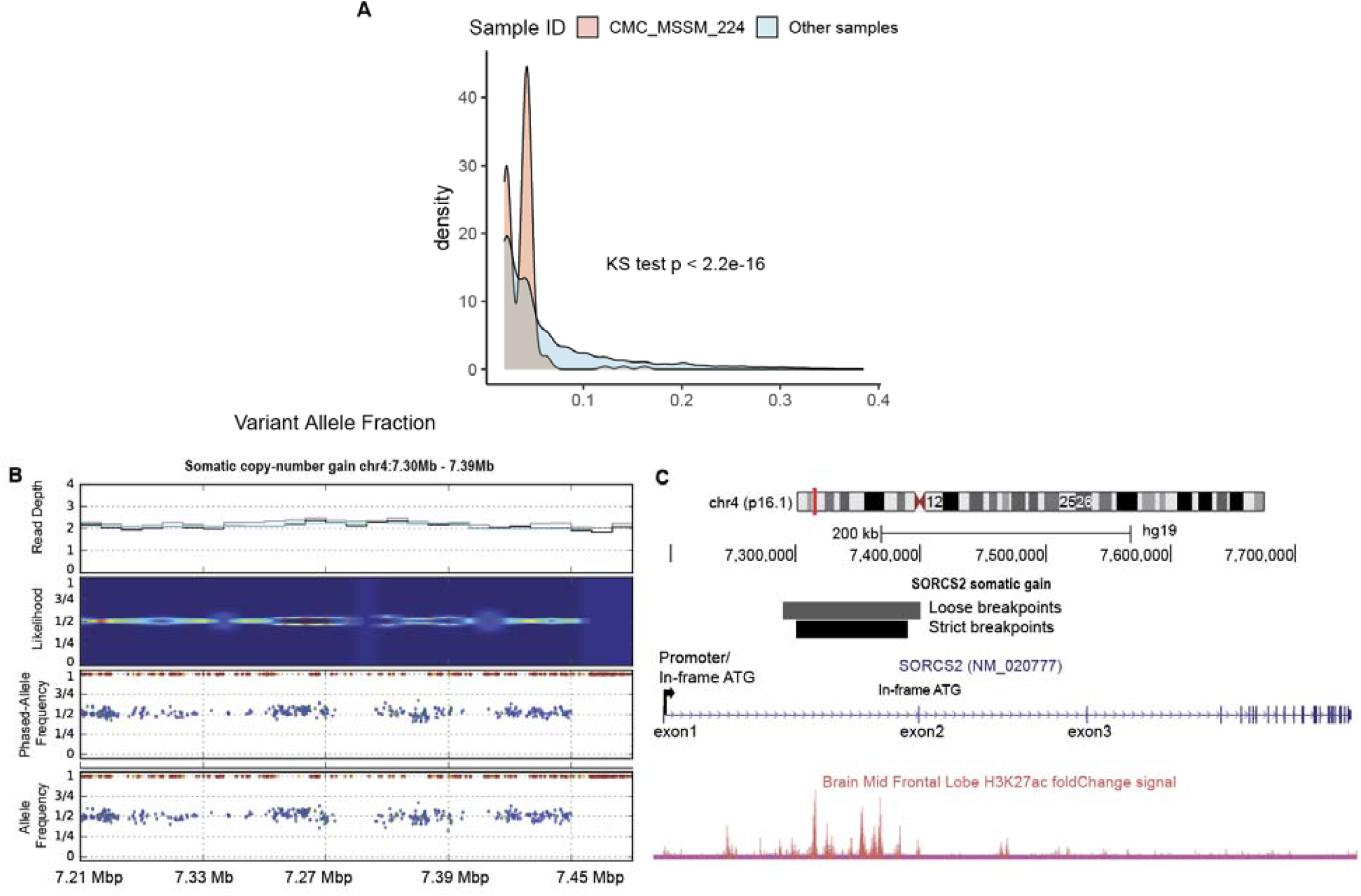
Variant allele fraction distribution of outlier sample and somatic gain in schizophrenia. A) Density plot of variant allele fraction in CMC_MSSM_224 sample compared to the rest of the samples. P-value was computed using the Kolmorogrov-Smirnov test. B) Plot of somatic gain in a SCZ sample. Heatmap represents the likelihood function with the red dotted lines indicating the likely event breakpoints. The dot plots indicate the phased and un-phased SNP allele-frequency respectively. C) A Genome Browser schematic of *SORCS2* gene with H3K27ac histone mark track from Roadmap Epigenomics consortium^32^.

**Figure S2.**
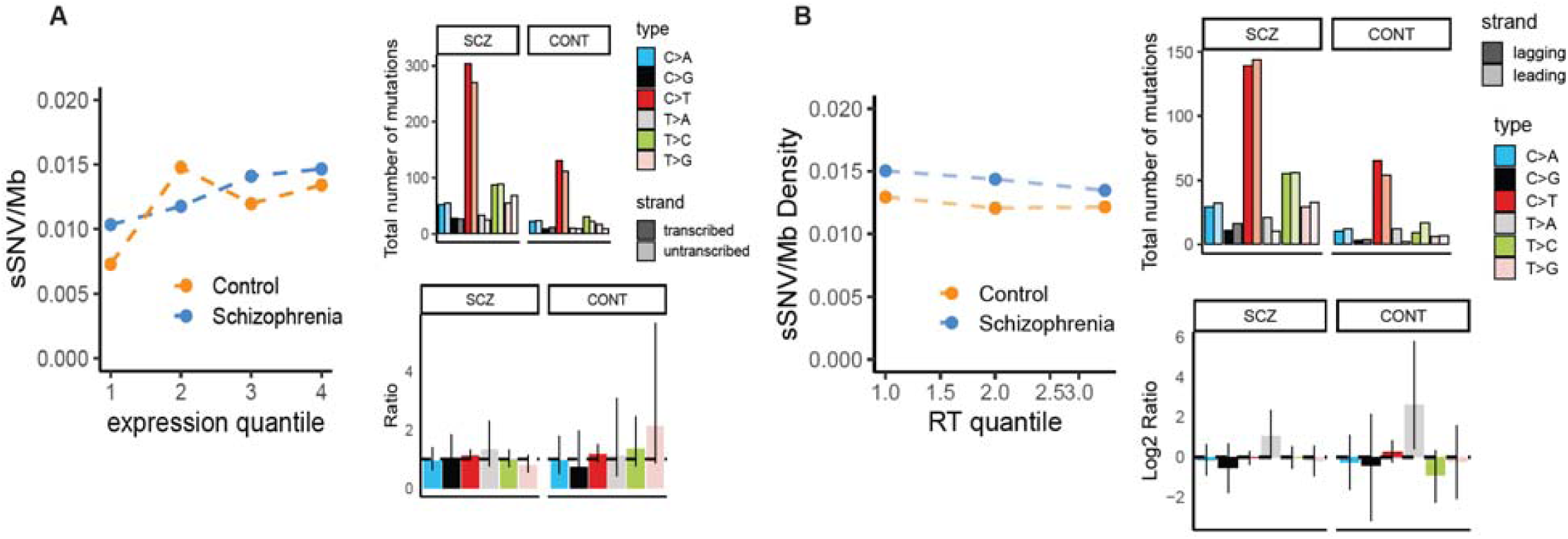
Characterization of transcriptional and replication strand bias in schizophrenia and controls. A) Scatter plot of fetal brain gene expression versus sSNV rate in schizophrenia cases and controls across expression quartiles. Bar plots showing genome-wide transcriptional strand bias as difference in total number of mutations and rate ratio. B) Scatter plot of replication time versus sSNV rate in schizophrenia cases and controls across terciles. Bar plots showing genome-wide replication strand bias as difference in total number of mutations and log2(rate ratio).

**Figure S3.**
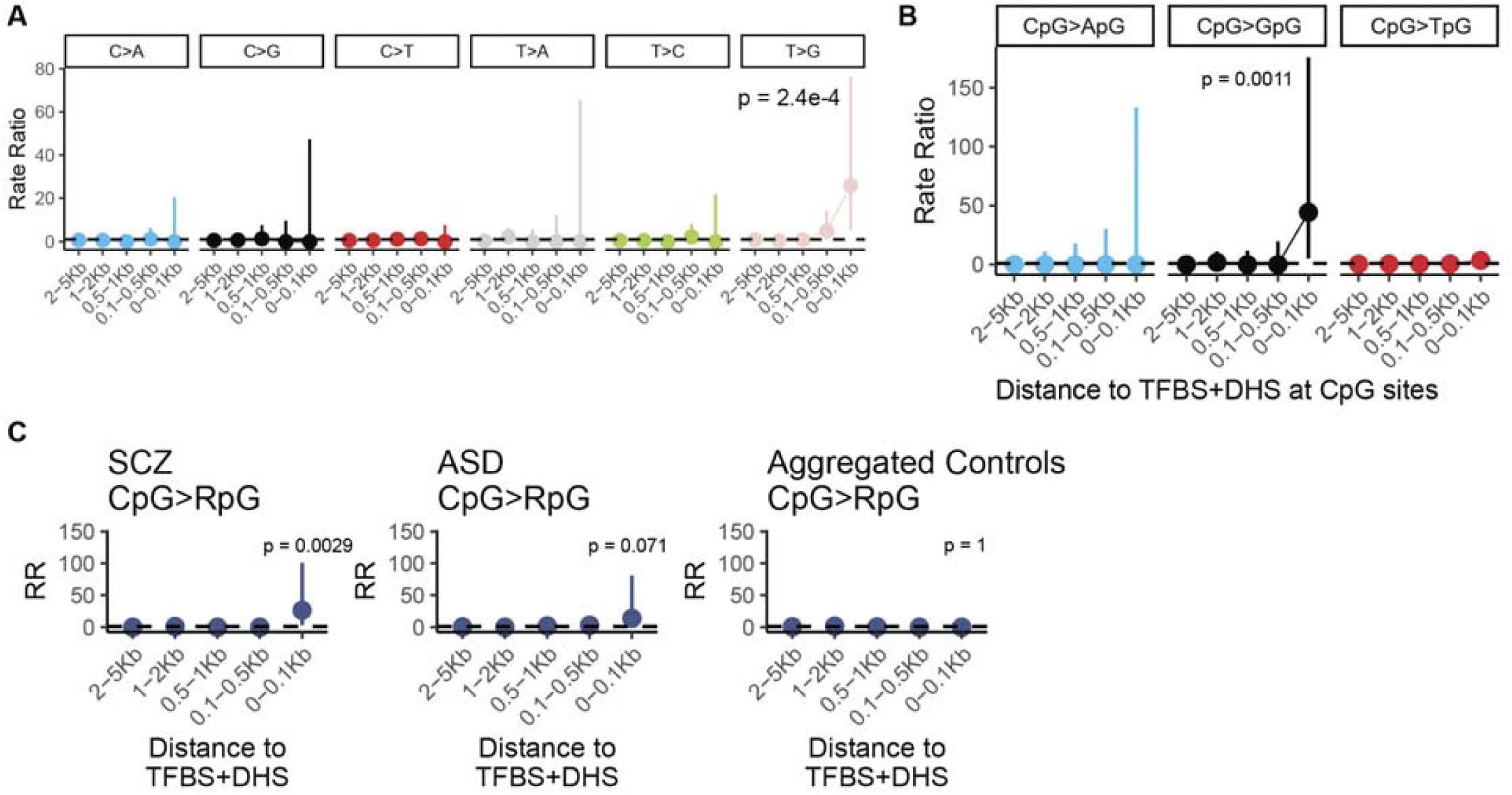
Relative sSNV rate around active TFBS. A) Forest plots of rate ratios in schizophrenia of different base changes in active TFBS at non-CpG sites and B) at CpG sites. C) Forest plots of rate ratios of CpG transversions active TFBS from schizophrenia, Autism Spectrum Disorder, and aggregated control sSNV. For panels A, B, and C p-values and 95% confidence intervals were computed using a Poisson test.

**Figure S4.**
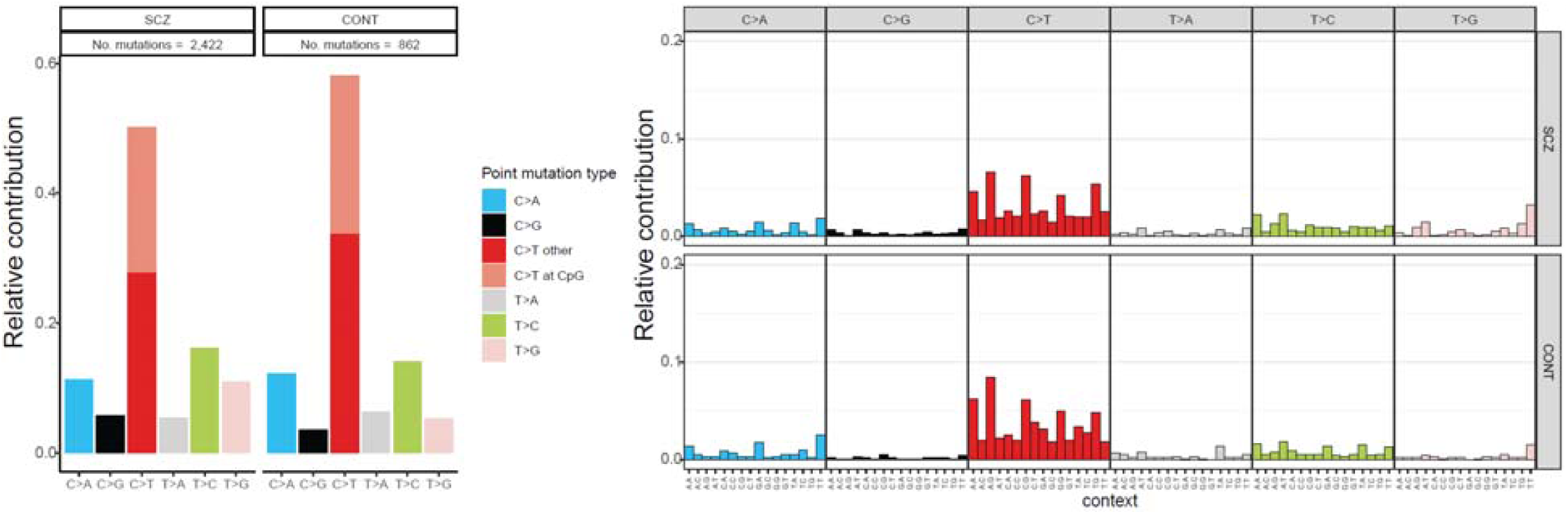
Single base substitution spectrum in schizophrenia and controls. (Left) Bar plot of single base substitution spectrum in schizophrenia case and controls normalized to the total number of sSNV in each diagnostic category. (Right) Bar plots of the tri-nucleotide context distribution in schizophrenia cases and controls, normalized to the total number of sSNV in each diagnostic category.

**Figure S5.**
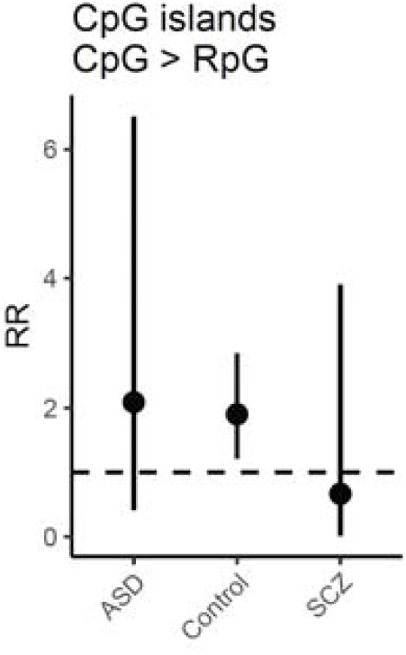
Relative somatic CpG transversion rate at CpG islands. Forest plot of the relative sSNV rate of CpG transversions at CpG islands compared to the genome-wide rate across diagnostic categories.

## Supplementary Data

**Table S1: BSMN member names and affiliations**

**Table S2: Subject clinical and demographic data**

**Table S3: Somatic mutation rates across transcription factor binding sites**

**Table S4: Recurrent somatic mutations**

**Table S5: DNAse hypersensitivity track descriptions**

## References

1. Singh, T., Neale, B. M. & Daly, M. J. Exome sequencing identifies rare coding variants in 10 genes which confer substantial risk for schizophrenia on behalf of the Schizophrenia Exome Meta-Analysis (SCHEMA) Consortium*. medRxiv 2020.09.18.20192815 (2020) doi:10.1101/2020.09.18.20192815.

2. Marshall, C. R. et al. Contribution of copy number variants to schizophrenia from a genome-wide study of 41,321 subjects. Nat. Genet. 49, 27–35 (2017).

3. Huo, Y., Li, S., Liu, J., Li, X. & Luo, X. J. Functional genomics reveal gene regulatory mechanisms underlying schizophrenia risk. Nat. Commun. 10, 1–19 (2019).

4. Consortium, S. W. G. of the P. G., Ripke, S., Walters, J. T. & O’Donovan, M. C. Mapping genomic loci prioritises genes and implicates synaptic biology in schizophrenia. medRxiv 2020.09.12.20192922 (2020) doi:10.1101/2020.09.12.20192922.

5. Brown, A. S. & Patterson, P. H. Maternal Infection and Schizophrenia: Implications for Prevention. Schizophr. Bull. 37, 284–290 (2011).

6. Khandaker, G. M., Zimbron, J., Lewis, G. & Jones, P. B. Prenatal maternal infection, neurodevelopment and adult schizophrenia: a systematic review of population-based studies. Psychol. Med. 43, 239–257 (2013).

7. Estes, M. L. & McAllister, A. K. Maternal immune activation: Implications for neuropsychiatric disorders. Science 353, 772–777 (2016).

8. Yim, Y. S. et al. Reversing behavioral abnormalities in mice exposed to maternalinflammation. Nature 549, 482 (2017).

9. Kim, S. et al. Maternal gut bacteria promote neurodevelopmental abnormalities in mouse offspring. Nature 549, 528–532 (2017).

10. Alexandrov, L. B. et al. The repertoire of mutational signatures in human cancer. Nature 578, 94–101 (2020).

11. Bae, T. et al. Different mutational rates and mechanisms in human cells at pregastrulation and neurogenesis. Science 359, 550–555 (2018).

12. Bizzotto, S. et al. Landmarks of human embryonic development inscribed in somatic mutations. Science 371, 1249–1253 (2021).

13. Heinzen, E. L. Somatic variants in epilepsy – advancing gene discovery and disease mechanisms. Current Opinion in Genetics and Development vol. 65 1–7 (2020).

14. Lim, E. T. et al. Rates, distribution and implications of postzygotic mosaic mutations in autism spectrum disorder. Nat. Neurosci. 20, 1217–1224 (2017).

15. Dou, Y. et al. Postzygotic single-nucleotide mosaicisms contribute to the etiology of autism spectrum disorder and autistic traits and the origin of mutations. Hum. Mutat. 38, 1002–1013 (2017).

16. Krupp, D. R. et al. Exonic Mosaic Mutations Contribute Risk for Autism Spectrum Disorder. Am. J. Hum. Genet. 101, 369–390 (2017).

17. Rodin, R. E. et al. The landscape of somatic mutation in cerebral cortex of autistic and neurotypical individuals revealed by ultra-deep whole-genome sequencing. Nat. Neurosci. 24, 176–185 (2021).

18. Ganz, J. et al. Rates and Patterns of Clonal Oncogenic Mutations in the Normal Human Brain. Cancer Discov. 12, 172–185 (2022).

19. Lee, J. H. Somatic mutations in disorders with disrupted brain connectivity. Exp. Mol. Med. 48, e239–9 (2016).

20. Shirley, M. D. et al. Sturge–Weber Syndrome and Port-Wine Stains Caused by Somatic Mutation in GNAQ. N. Engl. J. Med. 368, 1971–1979 (2013).

21. Rodin, R. E. & Walsh, C. A. Somatic Mutation in Pediatric Neurological Diseases. Pediatr. Neurol. 87, 20–22 (2018).

22. D’Gama, A. M. et al. Somatic Mutations Activating the mTOR Pathway in Dorsal Telencephalic Progenitors Cause a Continuum of Cortical Dysplasias. Cell Rep. 21, 3754–3766 (2017).

23. Mirzaa, G. M. et al. Association of MTOR mutations with developmental brain disorders, including megalencephaly, focal cortical dysplasia, and pigmentary mosaicism. JAMA Neurol. 73, 836–845 (2016).

24. Møller, R. S. et al. Germline and somatic mutations in the MTOR gene in focal cortical dysplasia and epilepsy. Neurol. Genet. 2, (2016).

25. Lim, J. S. et al. Brain somatic mutations in MTOR cause focal cortical dysplasia type II leading to intractable epilepsy. Nat. Med. 21, 395–400 (2015).

26. Matevossian, A. & Akbarian, S. Neuronal nuclei isolation from human postmortem brain tissue. J. Vis. Exp. (2008) doi:10.3791/914.

27. Wang, Y. et al. Comprehensive identification of somatic nucleotide variants in human brain tissue. Genome Biol. 22, 92 (2021).

28. Lesch, K. P. et al. Molecular genetics of adult ADHD: Converging evidence from genome-wide association and extended pedigree linkage studies. J. Neural Transm. 115, 1573–1585 (2008).

29. Baum, A. E. et al. A genome-wide association study implicates diacylglycerol kinase eta (DGKH) and several other genes in the etiology of bipolar disorder. Mol. Psychiatry 13, 197–207 (2008).

30. Ollila, H. et al. Findings from bipolar disorder genome-wide association studies replicate in a Finnish bipolar family-cohort. Mol. Psychiatry 14, 351 (2009).

31. Lachman, H. M. Copy variations in schizophrenia and bipolar disorder. Cytogenetic and Genome Research vol. 123 27–35 (2009).

32. Roadmap Epigenomics Consortium et al. Integrative analysis of 111 reference human epigenomes. Nature 518, 317–329 (2015).

33. Cai, Y. et al. H3K27me3-rich genomic regions can function as silencers to repress gene expression via chromatin interactions. Nat. Commun. 12, 1–22 (2021).

34. Sabarinathan, R., Mularoni, L., Deu-Pons, J., Gonzalez-Perez, A. & Lopez-Bigas, N. Nucleotide excision repair is impaired by binding of transcription factors to DNA. Nature 532, 264–267 (2016).

35. Frigola, J., Sabarinathan, R., Gonzalez-Perez, A. & Lopez-Bigas, N. Variable interplay of UV-induced DNA damage and repair at transcription factor binding sites. Nucleic Acids Res. 49, 891–901 (2021).

36. Katainen, R. et al. CTCF/cohesin-binding sites are frequently mutated in cancer. Nat. Genet. 47, 818–821 (2015).

37. Perera, D. et al. Differential DNA repair underlies mutation hotspots at active promoters in cancer genomes. Nature 532, 259–263 (2016).

38. Marin, M., Karis, A., Visser, P., Grosveld, F. & Philipsen, S. Transcription Factor Sp1 Is Essential for Early Embryonic Development but Dispensable for Cell Growth and Differentiation. Cell 89, 619–628 (1997).

39. Mascrez, B., Ghyselinck, N. B., Chambon, P. & Mark, M. A transcriptionally silent RXRα supports early embryonic morphogenesis and heart development. Proc. Natl. Acad. Sci. 106, 4272–4277 (2009).

40. Timberlake, A. T. et al. Co-occurrence of frameshift mutations in SMAD6 and TCF12 in a child with complex craniosynostosis. Hum. Genome Var. 5, 1–5 (2018).

41. Velazquez, F. N. et al. Brain development is impaired in c-fos -/- mice. Oncotarget 6, 16883 (2015).

42. Mao, P. et al. ETS transcription factors induce a unique UV damage signature that drives recurrent mutagenesis in melanoma. Nat. Commun. 9, 1–13 (2018).

43. Roberts, J. D. et al. An International Multicenter Evaluation of Type 5 Long QT Syndrome: A Low Penetrant Primary Arrhythmic Condition. Circulation 429–439 (2020) doi:10.1161/CIRCULATIONAHA.119.043114.

44. Seplyarskiy, V. B. et al. Population sequencing data reveal a compendium of mutational processes in the human germ line. Science 373, 1030–1035 (2021).

45. Wu, X. & Zhang, Y. TET-mediated active DNA demethylation: mechanism, function and beyond. Nat. Rev. Genet. 18, 517–534 (2017).

46. Chan, K., Resnick, M. A. & Gordenin, D. A. The choice of nucleotide inserted opposite abasic sites formed within chromosomal DNA reveals the polymerase activities participating in translesion DNA synthesis. DNA Repair (Amst). 12, 878–889 (2013).

47. Jonsson, H. et al. Differences between germline genomes of monozygotic twins. Nat. Genet. 53, 27–34 (2021).

48. Silbereis, J. C., Pochareddy, S., Zhu, Y., Li, M. & Sestan, N. The Cellular and Molecular Landscapes of the Developing Human Central Nervous System. Neuron 89, 248–268 (2016).

49. Eckersley-Maslin, M. A., Alda-Catalinas, C. & Reik, W. Dynamics of the epigenetic landscape during the maternal-to-zygotic transition. Nat. Rev. Mol. Cell Biol. 2018 197 19, 436–450 (2018).

50. Wang, J. et al. Persistence of RNA transcription during DNA replication delays duplication of transcription start sites until G2/M. Cell Rep. 34, 108759 (2021).

51. Li, H. & Durbin, R. Fast and accurate short read alignment with Burrows-Wheeler transform. Bioinformatics 25, 1754–1760 (2009).

52. Poplin, R. et al. Scaling accurate genetic variant discovery to tens of thousands of samples. bioRxiv 201178 (2017) doi:10.1101/201178.

53. Karczewski, K. J. et al. Variation across 141,456 human exomes and genomes reveals the spectrum of loss-of-function intolerance across human protein-coding genes. bioRxiv 531210 (2019) doi:10.1101/531210.

54. Suvakov, M., Panda, A., Diesh, C., Holmes, I. & Abyzov, A. CNVpytor: a tool for copy number variation detection and analysis from read depth and allele imbalance in whole-genome sequencing. Gigascience 10, 1–9 (2021).

55. Untergasser, A. et al. Primer3—new capabilities and interfaces. Nucleic Acids Res. 40, e115–e115 (2012).

56. Koressaar, T. & Remm, M. Enhancements and modifications of primer design program Primer3. Bioinformatics 23, 1289–1291 (2007).

57. Vorontsov, I. E. et al. Genome-wide map of human and mouse transcription factor binding sites aggregated from ChIP-Seq data 06 Biological Sciences 0604 Genetics. BMC Res. Notes 11, 1–3 (2018).

58. Funk, C. C. et al. Atlas of Transcription Factor Binding Sites from ENCODE DNase Hypersensitivity Data across 27 Tissue Types. Cell Rep. 32, (2020).

59. Girskis, K. M. et al. Rewiring of human neurodevelopmental gene regulatory programs by human accelerated regions. Neuron 109, 3239-3251.e7 (2021).

60. Akalin, A., Franke, V., Vlahoviček, K., Mason, C. E. & Schübeler, D. genomation: a toolkit to summarize, annotate and visualize genomic intervals. Bioinformatics 31, 1127–1129 (2015).

61. Blokzijl, F., Janssen, R., van Boxtel, R. & Cuppen, E. MutationalPatterns: Comprehensive genome-wide analysis of mutational processes. Genome Med. 10, 1–11 (2018).

