## Supplemental Text for "Enrichment of somatic mutations in schizophrenia brain targets prenatally active transcription factor bindings sites"

### Outlier SCZ sample

We noted that there was one outlier SCZ sample with 188 mutations. We could not identify any quality metrics, technical or artifactual features that would explain why this one sample showed more mutations than the rest. We did find that the VAF distribution of this sample showed a distinctive bimodal distribution (Figure S1A) probably reflecting a clonal expansion event that occurred prenatally, which has been reported to occur in non-diseased brains[1], thus we retained that sample for all downstream analyses. The excess in genome-wide burden in cases vs controls remained significant even after excluding this sample (p=7.76e-5).

### Somatic copy number gain in SCZ sample

Somatic CNV (sCNV) calling of our samples revealed a somatic gain in one SCZ sample and none in the control samples. We applied CNVpytor[2], a method developed to specifically identify sCNVs from high coverage WGS data on our case and control samples (see Methods). With this approach we observed one high confidence ~90Kb somatic gain mapping to chromosome 4 in a SCZ sample (Figure S1B,C). The somatic gain had an estimated 17.4% mosaic fraction and overlapped intron 1 of the *SORCS2* gene and possible exon2 (Figure S1B,C). We attempted to look for split reads supporting the exact breakpoints. Given our ~200x coverage we expected ~16 supporting reads, but were not able to identify them. However, upon closer inspection we found 2 simple repeat regions at both ends of the estimated breakpoints, hindering mappability at the breakpoint loci, thus it is not surprising we were not able to find supportive reads. Nevertheless, the shift in phased-allele frequency provides strong statistical evidence of an event in this region (Figure S1B,C). The simple repeat regions suggests that this sCNV potentially arose through tandem duplication. Thus, we expanded the range by 10 Kb around the estimated breakpoints to provide less stringent breakpoints (Figure S1B,C).

*SORCS2* encodes for a subunit of the sortilin-related VPS10 domain-containing receptor proteins, which are cell-surface proteins implicated in central nervous system development [3]. Previous GWAS and germline CNV studies have shown that variants in the *SORCS2* locus may confer risk for attention-deficit hyperactive disorder (ADHD), and bipolar disorder [4–7]. Similarly, germline SNPs in intron 1 of *SORCS2* have been associated with clinical outcomes in ADHD [8], suggesting a potential role in neuropsychiatric disease. Our somatic gain overlaps H3K27ac regions present in frontal cortex brain (Figure S1B,C), suggesting a potential dysregulation of the expression of this gene by altering enhancer interactions. If the breakpoints actually disrupt exon2, a frame-shift might result in an aberrant protein. *SORCS2* deficient mice display behavioral changes such as long-term memory formation impairment, increased risk taking, and ADHD-like behavior, and social memory deficits [9–12]. However, whether somatic gains within intron 1/exon2 of the *SORCS2* gene plays a role in SCZ requires further functional studies.

### References

1. Ganz, J. *et al.* Rates and Patterns of Clonal Oncogenic Mutations in the Normal Human

Brain. *Cancer Discov.* **12**, 172–185 (2022).

2. Suvakov, M., Panda, A., Diesh, C., Holmes, I. & Abyzov, A. CNVpytor: a tool for copy number variation detection and analysis from read depth and allele imbalance in whole-genome sequencing. *Gigascience* **10**, 1–9 (2021).

3. Hermey, G. *et al.* The three sorCS genes are differentially expressed and regulated by synaptic activity. *J. Neurochem.* **88**, 1470–1476 (2004).

4. Lesch, K. P. *et al.* Molecular genetics of adult ADHD: Converging evidence from genome-wide association and extended pedigree linkage studies. *J. Neural Transm.* **115**, 1573–1585 (2008).

5. Baum, A. E. *et al.* A genome-wide association study implicates diacylglycerol kinase eta (DGKH) and several other genes in the etiology of bipolar disorder. *Mol. Psychiatry* **13**, 197–207 (2008).

6. Ollila, H. *et al.* Findings from bipolar disorder genome-wide association studies replicate in a Finnish bipolar family-cohort. *Mol. Psychiatry* **14**, 351 (2009).

7. Lachman, H. M. Copy variations in schizophrenia and bipolar disorder. *Cytogenetic and Genome Research* vol. 123 27–35 (2009).

8. Alemany, S. *et al.* New suggestive genetic loci and biological pathways for attention function in adult attention-deficit/hyperactivity disorder. *Am. J. Med. Genet. Part B Neuropsychiatr. Genet.* **168**, 459–470 (2015).

9. Glerup, S. *et al.* SorCS2 regulates dopaminergic wiring and is processed into an apoptotic two-chain receptor in peripheral glia. *Neuron* **82**, 1074–1087 (2014).

10. Glerup, S. *et al.* SorCS2 is required for BDNF-dependent plasticity in the hippocampus. *Mol. Psychiatry* **21**, 1740–1751 (2016).

11. Olsen, D. *et al.* Altered dopaminergic firing pattern and novelty response underlie ADHD-like behavior of SorCS2-deficient mice. *Transl. Psychiatry* **11**, 1–14 (2021).

12. Yang, J. *et al.* SorCS2 is required for social memory and trafficking of the NMDA receptor. *Mol. Psychiatry* **26**, 927–940 (2021).
